# Graph topology reframes the coherence of cell-state manifold inference under heterogeneous single-cell observations

**DOI:** 10.64898/2025.12.13.694096

**Authors:** Tomohiro Tamura, Yusuke Yamane, Yuji Okano, Tetsuo Ishikawa, Kazuhiro Sakurada

**Author notes:** Corresponding author; Keio University School of Medicine and Graduate School of Medicine; Shinanomachi 35, Shinjuku, Tokyo, Japan, 160-8582. These authors equally contributed to this work (Tomohiro Tamura, Yusuke Yamane).

## Abstract

Manifold-based single-cell omics analyses assume that high-dimensional observations can fall into a low-dimensional space encoding biological constraints. In practice, per-cell observation is highly heterogeneous: shallowly- and deeply-observed cells coexist. In an empirical scRNA-seq dataset, shallowly-observed cells cluster together to generate spurious hubs that can give rise to illusory loops in low-dimensional manifold skeletons in graph abstraction. Several imputation methods leave these artifacts largely intact, whereas graph abstraction restricted to homogeneously-observed cells alone recovers tree-like structures locally representing constrained cell state transitions. Simulations further demonstrate that realistic heterogeneous observation can create spurious subclusters and false branching. Grounded by these observations, we propose topological stability descriptors of low-dimensional manifold skeletons to delineate a regime in which manifold-based inference is trustworthy despite realistic heterogeneous observations. Our findings underscore that the heterogeneity of observations is not merely noise but a source of systemic distortion in manifold-based inference that must be addressed.

## Introduction

Manifold-based analyses have become a central paradigm in single-cell transcriptomics^1–3^. In this view, each cell is represented as a point in a high-dimensional data space, but observations of biologically attainable states are assumed to lie near a low-dimensional manifold that encodes constraints such as differentiation paths and responses to stimuli. By examining local neighborhood relations in this space and constructing graphs to abstract cellular state similarity, one hopes to recover the coarse-grained, low-dimensional structure from snapshot data and to interpret it as a trajectory of state transitions.

This paradigm implicitly relies on an ideal regime of measurement in which cellular states are represented by uniformly informative observations. In practice, however, droplet-based scRNA-seq produces datasets with pronounced heterogeneous observation: shallowly- and deeply observed cells coexist, so that some cells possess rich molecular profiles to represent their states whereas others appear to have only sparse snapshots. Prior work has characterized how technical noise and gene-level *dropout* affect dimensionality reduction and clustering, and a wide range of imputation and normalization methods has been proposed and discussed to mitigate these effects^4–6^. Much less is known, however, about how heterogeneous single-cell observations themselves affects the geometry and topology of inferred data manifolds, and under what conditions manifold-based inferences remain reliable when shallow and deep observations are mixed.

Here we first show how realistic heterogeneous observations can systematically distort manifold-based inference. Using a peripheral blood mononuclear cell (PBMC) dataset, we demonstrate that shallowly-observed cells in heterogeneous observations form spurious hubs and illusory loops in graph abstraction, whereas restricting analyses to homogeneously-observed cells recovers a tree-like structure locally representing constrained cell state transitions. Computational simulations reveal how realistic levels of heterogeneous observations can give rise to spurious subclusters, artifactual intermediates, and false branching. Motivated by these observations, we propose topological stability descriptors of low-dimensional manifold skeletons to delineate a regime in which manifold-based inference is trustworthy despite realistic heterogeneous observations. The unavoidable heterogeneity of observations is not merely noise but a major source of systemic distortion in manifold-based inference that must be addressed.

## Results

### Graph abstraction from heterogeneous single-cell observations generates a loop-rich structure

In general, scRNA-seq data are analyzed following the workflow, which we denote (#): log1p normalization, identification of highly variable genes (HVGs), scaling of gene expression across cells, principal component analysis (PCA), neighbor-graph construction, unsupervised clustering, and non-linear dimensionality regression uniform manifold approximation and projection (UMAP). To investigate whether heterogeneous observations empirically influences manifold-based inference in scRNA-seq data, we first applied the workflow (#) to viable cells in a PBMC scRNA-seq dataset^7^. As a result, Louvain clustering broadly classified cells into monocyte populations (clusters 0, 4, 5, 10, 12, 13, and 17), lymphocyte populations, and other cell types such as platelets (Figure 1A, Supplementary Figure 1A). When total unique molecular identifier (UMI) counts were projected onto the z-axis of these UMAP embeddings, shallowly-observed cells within monocyte populations were confined to specific regions (Supplementary Figure 1B). To investigate this biased distribution more closely, we subsequently reanalyzed the seven clusters and defined a putative monocyte population *M* (4,410 cells) by removing small cell populations representing technical artifacts (Supplementary Figure 1C, D). Applying the workflow (#) and the three-dimensional visualization to *M* recapitulated the biased distribution of shallowly-observed cells in the manifold-based inference, as observed in the global analysis (Figure 1B–D, Supplementary Figure 1E). Notably, cluster 1 was predominantly composed of shallowly-observed cells, and gave rise to a pronounced heavy tail in the distribution of per-cell total UMI counts. Moreover, application of the workflow (#) to a putatively tissue-derived cell population in a mouse scRNA-seq dataset from a benchmark study^8^ likewise identified an aggregation of shallowly observed cells in UMAP, suggesting that this is not a dataset-specific phenomenon (Supplementary Figure 1F, G).

**Figure 1.**
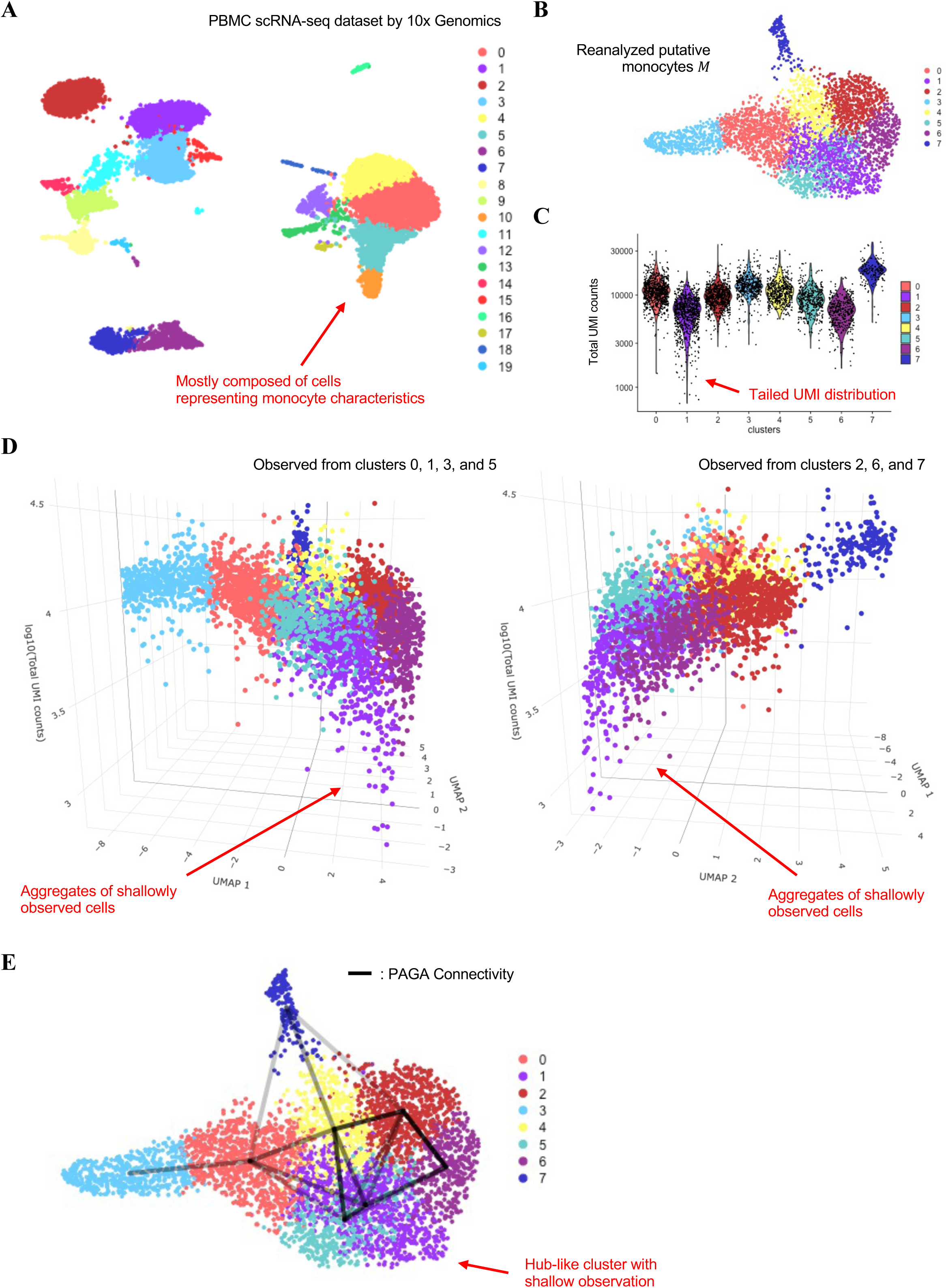
Low-information cells confined to specific regions form spurious hub-like connectivity in graph-based abstraction. **A** For viable cells with < 15% mitochondrial RNA content in a public PBMC scRNA-seq dataset by 10x Genomics, we applied a standard analysis workflow (#) consisting of log1p normalization, highly variable gene (HVG) selection, scaling, principal component analysis (PCA), nearest-neighbor graph construction, Louvain clustering, and UMAP embedding. The UMAP plot is shown with the resulting cluster annotations. Among these, clusters 0, 4, 5, 10, 12, 13, and 17 (red arrows) exhibit monocyte-like gene expression profiles and form a continuous distribution in UMAP space. The remaining clusters correspond to plausible peripheral blood populations such as lymphocytes and platelets; **B** After removing samples suspected of representing technical contamination by mRNA derived from multiple cells, we re-applied the workflow (#) to the putative monocyte population *M*. The UMAP plot is shown with the resulting cluster annotations; **C** Violin plot of total UMI counts per cell in each cluster. Cluster 1 (red arrow) is particularly enriched for shallowly-observed cells and exhibits a characteristic heavy-tailed distribution; **D** Dot plots visualizing the three-dimensional distribution of the putative monocyte population, with UMAP coordinates on the x–y plane and total UMI counts on the z-axis. Panels show views highlighting clusters 0, 1, and 3, and clusters 2, 4, 6, and 7, respectively. Shallowly-observed cells in cluster 1 (red arrow) are particularly confined to a specific region of the UMAP manifold; **E** UMAP embedding of *M* overlaid with partition-based graph abstraction (PAGA) edges representing inter-cluster connectivity. Only edges with connectivity ≥ 0.05 are shown, and the intensity of each line segment reflects the strength of connectivity. Cluster 1 (red arrow), which is predominantly composed of shallowly-observed cells, shows strong connectivity with many other clusters; Interactive HTML-based visualization tools for all three-dimensional plots are provided as **supplementary data**.

We next inferred PC neighbor graph-based connectivity among the clusters within *M* that were coarsely identified as distinct states using partition-based graph abstraction (PAGA)^9^. When we drew edges on the UMAP embedding between cluster pairs whose PAGA-inferred connectivity exceeded a given threshold (e.g., ≥ 0.05), a loop-rich structure emerged among clusters (Figure 1E). Interpreting this graph structure as a simplicial complex and analyzing it topologically revealed nine independent loops; particularly, the cluster 1, prominently composed of shallowly-observed cells, was connected by edges to clusters 0, 2, 4, 5, and 6, and thus, at face value, would be interpreted as a hub-like cell population in manifold-based inference.

### Graph abstraction using homogeneous observation alone generates a low-dimensional skeleton locally representing constrained cell state transitions

To investigate whether the results of graph abstraction behave differently without extremely heterogeneous observations containing shallowly-observed cells which formed the spurious hub-like subclusters, we extracted 1,986 cells with total UMI counts > 10,000 from *M* to define a subset *M*′, and applied the workflow (#) to *M*′. As a result, cells were partitioned into seven clusters, and any cluster with the pronounced heavy tail in the distribution of per-cell total UMI counts observed in *M* was no longer apparent (Figure 2A, Supplementary Figure 2A, B). Connectivity inference using PAGA further revealed a simpler, tree-like structure centered around cluster 2 as a hub (Figure 2B). Moreover, differentially expressed gene analyses enabled biologically interpretable annotation of each cluster, capturing several plausible local structures of monocyte state transitions consistent with prior experimental and clinical work as follow (Figure 2C–G, Supplementary Figure 2C):

**Figure 2.**
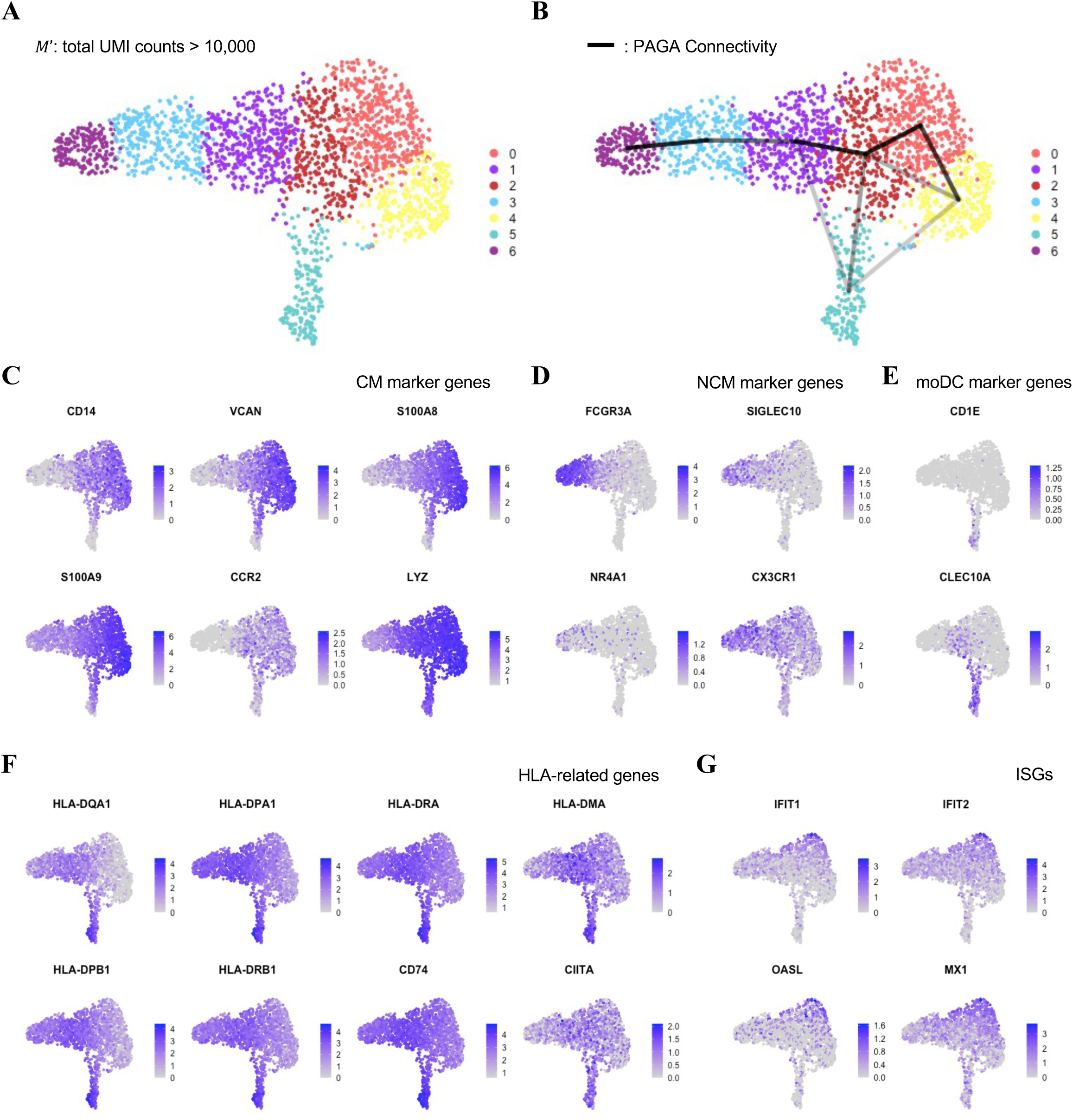
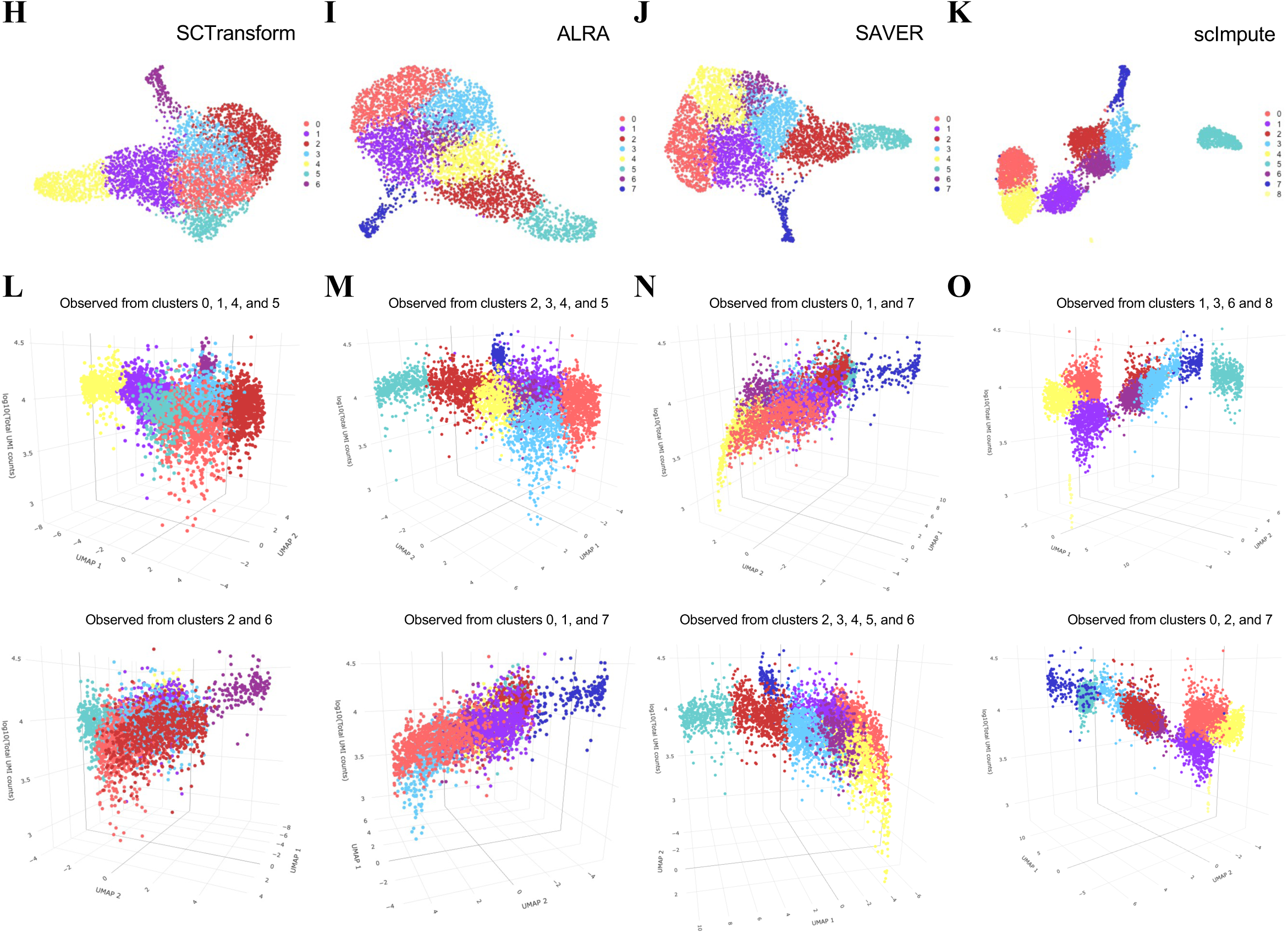
Graph-based data abstraction using high-information cells alone generates plausible inference of cell state transition. **A** From the putative monocyte population *M*, we extracted high-information cells defined by total UMI counts ≥ 10,000 to obtain subset *M*′, and applied the workflow (#) to *M*′. The UMAP plot is shown with the resulting cluster annotations; **B** UMAP embedding of *M*′ overlaid with PAGA edges representing inter-cluster connectivity. Only edges with connectivity ≥ 0.05 are shown, and the intensity of each line segment reflects the strength of connectivity; **C** Feature plots showing the expression of classical monocyte (CM) marker genes *CD14*, *VCAN*, *S100A8*, *S100A9*, *CCR2*, and *LYZ* in *M*′. The color bar indicates normalized expression levels; **D** Feature plots showing the expression of non-classical monocyte (NCM) marker genes *FCGR3A*, *SIGLEC10*, *NR4A1*, and *CX3CR1* in *M*′. The color bar indicates normalized expression levels; **E** Feature plots showing the expression of monocyte-derived dendritic cell (moDC) marker genes *CD1E* and *CLEC10A* in *M*′ . The color bar indicates normalized expression levels; **F** Feature plots showing the expression of HLA-related genes *HLA-DPA1*, *HLA-DQA1*, *HLA-DRA*, *HLA-DMA*, *HLA-DPB1*, *HLA-DRB1*, *CD74*, and *CIITA* in *M*′. The color bar indicates normalized expression levels; **G** Feature plots showing the expression of IFN-stimulated genes (ISGs) *IFIT1*, *IFIT2*, *OASL*, and *MX1* in *M*′. The color bar indicates normalized expression levels; **H** After normalizing gene expression profiles in *M* using SCTransform, we applied the workflow (#). The UMAP plot is shown with the resulting cluster annotations; **I** After imputing *dropouts* in *M* using ALRA, we applied the workflow (#). The UMAP plot is shown with the resulting cluster annotations; **J** After imputing *dropouts* in *M* using SAVER, we applied the workflow (#). The UMAP plot is shown with the resulting cluster annotations; **K** After imputing *dropouts* in *M* using scImpute with a parameter *k* = 7 which is the same number of clusters as stratified in *M*′, we applied the workflow (#). The UMAP plot is shown with the resulting cluster annotations; **L** Dot plots visualizing the three-dimensional distribution of the SCTransform-normalized *M*, with UMAP coordinates on the x–y plane and total UMI counts on the z-axis; **M** Dot plots visualizing the three-dimensional distribution of the ALRA-imputed *M*, with UMAP coordinates on the x–y plane and total UMI counts on the z-axis; **N** Dot plots visualizing the three-dimensional distribution of the SAVER-imputed *M*, with UMAP coordinates on the x–y plane and total UMI counts on the z-axis; **O** Dot plots visualizing the three-dimensional distribution of the scImpute-imputed *M*, with UMAP coordinates on the x–y plane and total UMI counts on the z-axis; Interactive HTML-based visualization tools for all three-dimensional plots are provided as **supplementary data**.

Annotations of each coarsely-discriminated cell state (CM: classical monocyte, IM: intermediate monocyte, NCM: non-classical monocyte, ISG: interferon (IFN)-stimulated gene, moDC: monocyte-derived dendritic cell):

- cluster 0: IFN-stimulated CMs (*HLA genes^low^*, *ISGs^high^*, *CM markers^high^*)
- cluster 1: IM-NCM-lineage-primed CMs (*HLA genes^high^*, *CM marker^high^*, *NCM markers^low^*)
- cluster 2: HLA^+^ basal CMs (*HLA genes^moderate^*, *CM markers^high^*)
- cluster 3: IMs (*HLA genes^high^*, *CM markers^moderate^*, *NCM markers^moderate^*)
- cluster 4: HLA^neg/low^ CMs potentially immunosuppressive or exhausted (*HLA genes^neg/low^*, *CM markers^high^*)
- cluster 5: NCMs (*HLA genes^moderate^*, *NCM markers^high^*)
- cluster 6: moDCs (*HLA genes^high^*, *DC markers^high^*) Local structures consistent with prior work^10–15^:
- connection 4−0−2: IFNγ-mediated HLA induction during monocyte maturation and immune regulation
- connection 2−1−3−5: Differentiation to NCMs from CMs with temporal HLA upregulation at IMs
- connection 2−6: Differentiation to moDCs from CMs

These results suggest that graph abstraction using homogeneously-observed cells alone can uncover a low-dimensional manifold skeleton locally representing constrained cell state transitions that was obscured by heterogeneous observations.

Uncertainty in scRNA-seq data arising from limited observation depth has been widely discussed in the context of *dropout*, and numerous methods have been developed to impute or normalize expression profiles for genes that are not detected due to technical limitations. We next examined how the realistic levels of heterogeneous observations affects such correction procedures and results. We applied four commonly used imputation or normalization approaches: SCTransform^16^, ALRA^17^, SAVER^18^, and scImpute^19^ for the *M*, and applied the workflow (#) to the imputed or normalized gene expression data (Figure 2H–K). As observed in the uncorrected *M* (Figure 1C, D), the three-dimensional visualizations with UMAP embedding and total UMI counts showed that shallowly-observed cells were consistently confined to specific regions that were captured as distinct Louvain clusters with the heavy tail of total UMI count distribution across the four methods (Figure 2L–O, Supplementary Figure 2D–K). These results indicate that the heterogeneity of per-cell observation is difficult to correct by *dropout* imputation, because it intrinsically induces gene-wise differences in the expected distribution of UMI counts across cells.

### Simulated cell population shows how heterogeneous observations generate spurious subclusters

To simulate the influences of heterogeneous observations on various interpretations generally applied to scRNA-seq data, we first modeled cDNA (UMI) intracellular distribution among genes using several empirical UMI count data^7,8,20^. We calculated *N_i_*(*k*), the number of genes with exactly *k* observed UMIs, for cell 𝑖 and computed *N̄*(*k*), the mean of *N_i_*(*k*), at each depth range (Supplementary Figure 3A–C). As a result, though dataset-specific differences were observed in the detailed behavior of *N̄*(*k*) such as a heavy- or light-tailed *N̄*(*k*) distribution, three common tendencies emerged:

1. *N̄*(*k*) monotonically decreased as *k* increased.
2. For 1 ≤ *k* ≤ 10, *N̄*(1) was much higher than those of the other *k*s, and *N̄*(*k*) decreased sharply.
3. *N̄*(*k*) decreased gradually for *k* > 10, especially in higher-depth ranges.

We next simulated gene-specific cDNA fractions in the cDNA pool that could reproduce the observed *N̄*(*k*) profiles under random sampling. For each gene, we denote its fraction in the cDNA pool as *p*, and assign gene index 𝑖 (𝑖 = 1, 2, . . . , 36,601) in ascending order of *p*. It has been estimated that the total number of genes expressed in a single cell is approximately 10,000; therefore, we assumed that only genes with indices 𝑖 ≥ 26,602 can be expressed^21,22^. In addition, the simulation was conducted under the assumption that all UMIs are equally probable and a distinct UMI is captured without replacements for each gene.

Consequently, we constructed three expression distribution models in which *p*_*i*_ increases as a function of 𝑖 in a linear, exponential, or combined manner (Figure 3A, Supplementary Figure 3D, E). Under Poisson or negative binomial (NB) sampling models with various dispersion parameters, we computed 𝔼[*N*(*k*)], expected *N*(*k*), and compared them with the empirical *N̄*(*k*), which indicated that the sharp decrease in *N̄*(*k*) for 1 ≤ *k* ≤ 10 and gradual decrease in *N̄*(*k*) for *k* > 10 could be simultaneously explained by the combined (both linear and exponential) increase of *p*_*i*_ (Figure 3B, C, Supplementary Figure 3F–I). From these results, we assumed that the expression distribution model, which we denote (*), with the combined increase of *p*_*i*_ at least could reproduce the UMI distribution similar to the empirical *N̄*(*k*), and overdispersion could underlie the differences in the detailed behaviors of *N̄*(*k*) across datasets.

**Figure 3.**
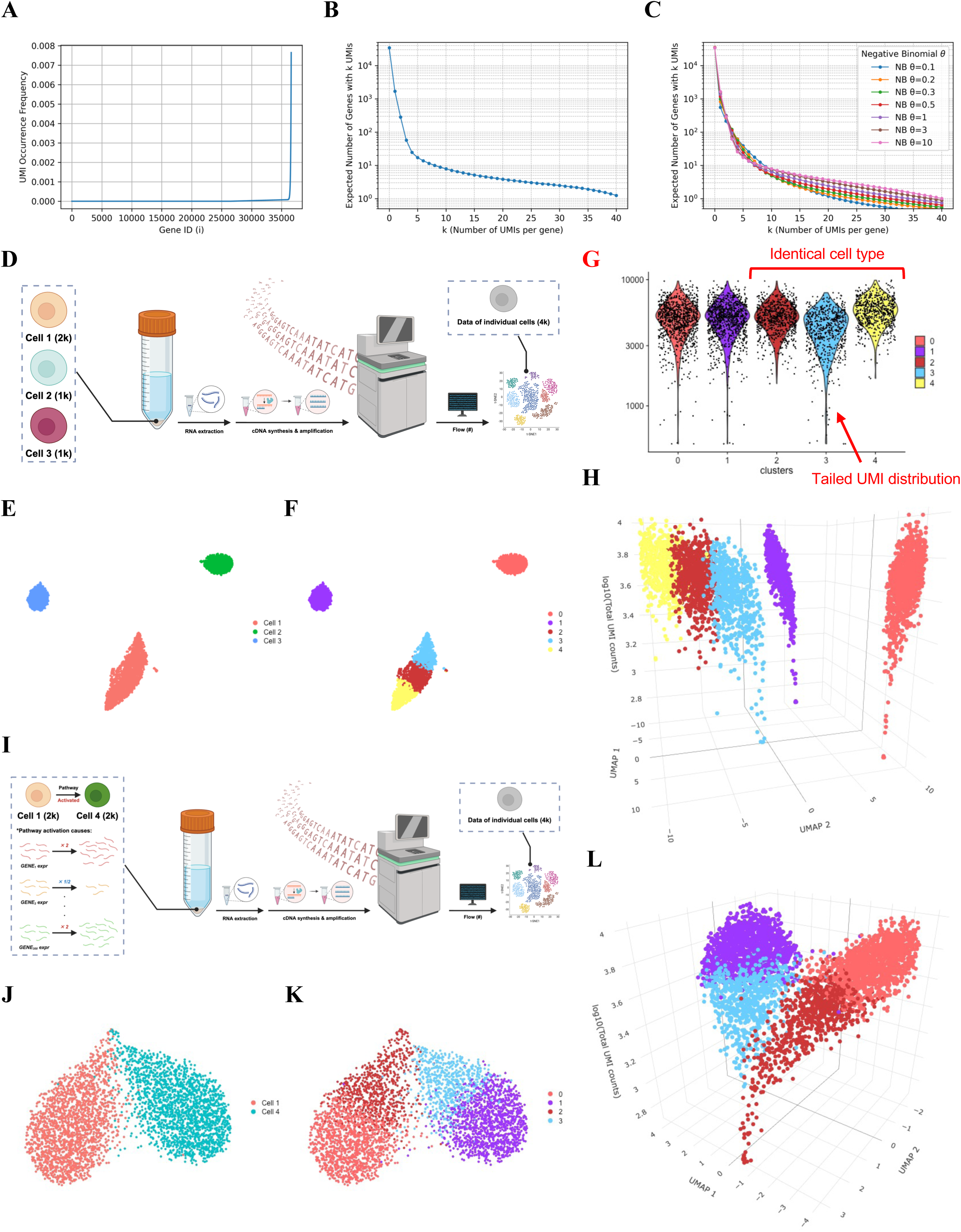
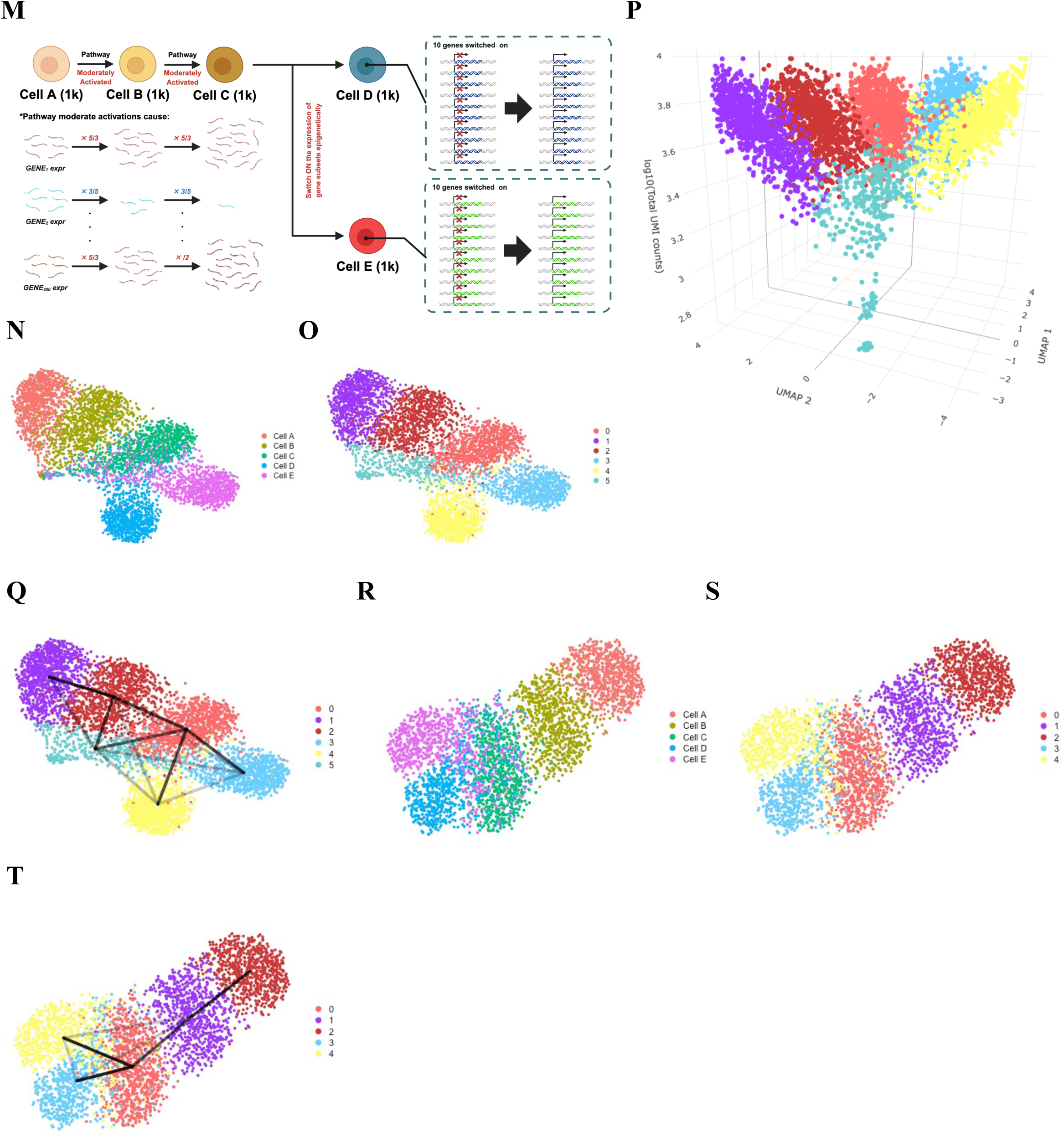
Simulations demonstrate how heterogeneous single-cell observation distorts cell-state discrimination and state trajectory inference. **A** A mixed (linear + exponential) distribution model for the composition ratio *p* of gene-derived products in the cDNA pool. The number of genes is set to 36,601, corresponding to a typical gene panel. For each gene, we model the molecule count as an integer so that the total number of molecules is on the order of 10^5^, and we use the values normalized by the total molecule count as *p*. Genes are indexed such that *p*_1_ ≤ *p*_2_ ≤ ⋯ ≤ *p*_36,601_. It is generally considered that approximately 10,000 genes are expressed in a single cell; in this model, gene expression is allowed only for 𝑖 ≥ 26,601. For 26,602 ≤ 𝑖 ≤ 36000, *p*_*i*_ increases linearly, and for 𝑖 ≥ 36,000, an additional term representing exponential increase is applied; **B** Line plot showing the expected number of genes *N*(*k*) with exactly *k* observed UMIs under a Poisson sampling model. For each gene, all UMIs are assumed to be sampled with equal probability, and a different UMI is selected at each draw; **C** Line plots showing the expected *N*(*k*) under a negative binomial sampling model. Expectations are computed for the overdispersion parameter *θ* = 0.1, 0.3, 0.5, 1, 3, and 10; **D** Schematic illustration of simulations modeling different cell-type discrimination in a simulated cell population 𝑋. Created with BioRender; **E** UMAP embedding of cell population 𝑋 after applying the workflow (#), annotated with ground-truth cell-type labels; **F** UMAP embedding of cell population 𝑋 after applying the workflow (#), annotated with clusters obtained by Louvain clustering; **G** Violin plots of total UMI counts per cell for each Louvain cluster in cell population 𝑋. Among clusters 3, 4, and 5, which are composed of observation samples from Cell 1, cluster 4 (red arrow) contains many low-information cells and exhibits a pronounced heavy-tailed distribution; **H** Dot plots visualizing the three-dimensional distribution of cell population 𝑋, with UMAP coordinates on the x–y plane and total UMI counts on the z-axis; **I** Schematic illustration of simulations modeling transcriptional changes (e.g., pathway activation) in a simulated cell population 𝑌. Created with BioRender; **J** UMAP embedding of cell population 𝑌 after applying the workflow (#), annotated with ground-truth cell-type labels; **K** UMAP embedding of cell population 𝑌 after applying the workflow (#), annotated with clusters obtained by Louvain clustering; **L** Dot plots visualizing the three-dimensional distribution of cell population 𝑌, with UMAP coordinates on the x–y plane and total UMI counts on the z-axis; **M** Schematic illustration of simulations modeling step-wise cell state transitions assuming pathway activation and epigenetic de-repression in a simulated cell population *Z*. Created with BioRender; **N** UMAP embedding of cell population *Z* after applying the workflow (#), annotated with ground-truth cell-type labels; **O** UMAP embedding of cell population *Z* after applying the workflow (#), annotated with clusters obtained by Louvain clustering; **P** Dot plots visualizing the three-dimensional distribution of cell population *Z*, with UMAP coordinates on the x–y plane and total UMI counts on the z-axis; **Q** UMAP embedding of *Z* overlaid with PAGA edges representing inter-cluster connectivity. Only edges with connectivity ≥ 0.05 are shown, and the intensity of each line segment reflects the strength of connectivity. Cluster 2 (red arrow), which is predominantly composed of shallowly-observed cells, shows strong connectivity with many other clusters; **R** From the putative monocyte population *Z*, we extracted high-information cells defined by total UMI counts ≥ 5,000 to obtain subset *Z*′, and applied the workflow (#) to *Z*′. The UMAP plot is shown with the ground-truth cell-type labels; **S** UMAP embedding of cell population *Z*′ after applying the workflow (#), annotated with clusters obtained by Louvain clustering; **T** UMAP embedding of *Z*′ overlaid with PAGA edges representing inter-cluster connectivity. Only edges with connectivity ≥ 0.05 are shown, and the intensity of each line segment reflects the strength of connectivity; Interactive HTML-based visualization tools for all three-dimensional plots are provided as **supplementary data**.

To evaluate how heterogeneous observations affect the generally used workflow (#), we first generated a simulated cell population 𝑋 consisting of 2,000 Cells 1 according to the model (*), and 1,000 Cells 2 and 3 modeling completely distinct cell-types from Cell 1 (Figure 3D). For each simulated cell, the total UMI counts were randomly drawn as an integer from a normal distribution with mean 5,000 and standard deviation 1,500, and clamped at a minimum of 500. When applying the workflow (#) to 𝑋, Cell 1, Cell 2, and Cell 3 were embedded as discrete groups in UMAP, but Louvain clustering interpreted spatial dispersion in Cell 1 as subclusters (Figure 3E, F). The three-dimensional visualization shows that the spatial dispersion in Cell 1 was associated with heterogeneous observation depth (Figure 3G, H, Supplementary Figure 3J).

We next generated a simulated cell population 𝑌 (Figure 3I) consisting of 2,000 Cells 1 and 2,000 Cells 4 modeling a pathway activation. When the workflow (#) was applied to 𝑌, Cell 1 and Cell 4 formed an apparently continuous distribution in UMAP, and Louvain clustering produced artifactual intermediates between Cell 1 and Cell 4 (Figure 3J, K). The three-dimensional visualization shows that distance in UMAP between Cell 1 and Cell 4 decreased as observations became shallow (Figure 3L, Supplementary Figure 3K). These results suggest that heterogeneous observations can generate spurious subclusters within certain cell-type populations or artifactual intermediates between subtypes.

### Simulated cell lineage shows how heterogeneous observations distort state transition inference

Such transcriptional changes simulated by 𝑌 are crucial for cellular differentiation and responses to environmental stimuli within tissues. Here, instead of assuming intrinsic continuous state transitions interpreted by *pseudotime*, we simulated step-wise cell state transitions and investigated the impact of heterogeneous observations on PAGA-inferred connectivity. To this end, we generated a simulated cell population *Z* with an assumed lineage: Cell A → Cell B (pathway activation) → Cell C (further pathway activation)→ Cell D or Cell E (epigenetic de-repression) (Figure 3M). We assume that there are 1,000 cells at each state.

When the workflow (#) and the three-dimensional visualization were applied to *Z*, the cells formed a continuous distribution in UMAP, and the distances among different states became progressively closer as observations became shallow, as observed in the previous simulations (Figure 3N, O). Louvain clustering yielded clusters in which shallowly-observed cells from different underlying states were intermixed (Figure 3P, Supplementary Figure 3L). When we generated connectivity graph between these clusters using PAGA, the intermixed clusters remained connected to multiple neighboring clusters, giving rise to an overly complex and artifactual loop-rich structure (Figure 3Q).

To assess whether the simple correction by enforcing a depth threshold can improve lineage inference, we generated *Z*′ by subsetting cells from *Z* with total UMI counts ≥ 4,000, and applied the workflow (#). In *Z*′, no spurious intermixed clusters were formed, and PAGA recovered strong connectivity specifically along the step-wise transition paths consistent with the assumed lineage (Figure 3R–T). These results are consistent with the results from the empirical PBMC scRNA-seq dataset and suggest the mechanism by which heterogeneous observations drive graph abstraction away from the manifold structure dictated by underlying biological constraints.

### Topology-guided diagnostics preserve the coherence of manifold inference while minimizing sample loss

Filtering for deeply-observed cells to generate a subset with homogeneous observations can uncover a low-dimensional manifold skeleton reflecting intrinsic paths of cell state transitions, but at the same time threatens the premise that each attainable state is sampled densely enough. Given prior reports that per-cell depth can influence PCA^23^, the plausibility of the latent space generated by the workflow (#) under heterogeneous observations needs to be examined explicitly. Moreover, when the workflow (#) is applied to all viable PBMCs, the distribution of total UMI counts vary substantially across cell types, indicating that although heterogeneous observations can distort manifold-based inference, simply thresholding cells by observation depth is biologically questionable because it ignores genuine differences in transcriptional activity (Supplementary Figure 4).

We defined cells included in *M*’ as high-information cells and the others filtered simply by observation depth as low-information cells, and evaluated PAGA connectivity among the annotated clusters in *M*’ (Figure 2A) and low-information cell population using the PCs learned on *M*. In this latent space, though the annotated clusters were connected with low-information cell population and generated numerous independent loops, the connectivity among the annotated clusters themselves was largely consistent with that observed when PCA was applied to *M*′ (Figure 2B, 4A). Indeed, when we applied downstream procedures of the workflow (#) to *M*′ using the PCs learned on *M*, the low-dimensional manifold skeleton was remarkably well preserved (Figure 4B).

**Figure 4.**
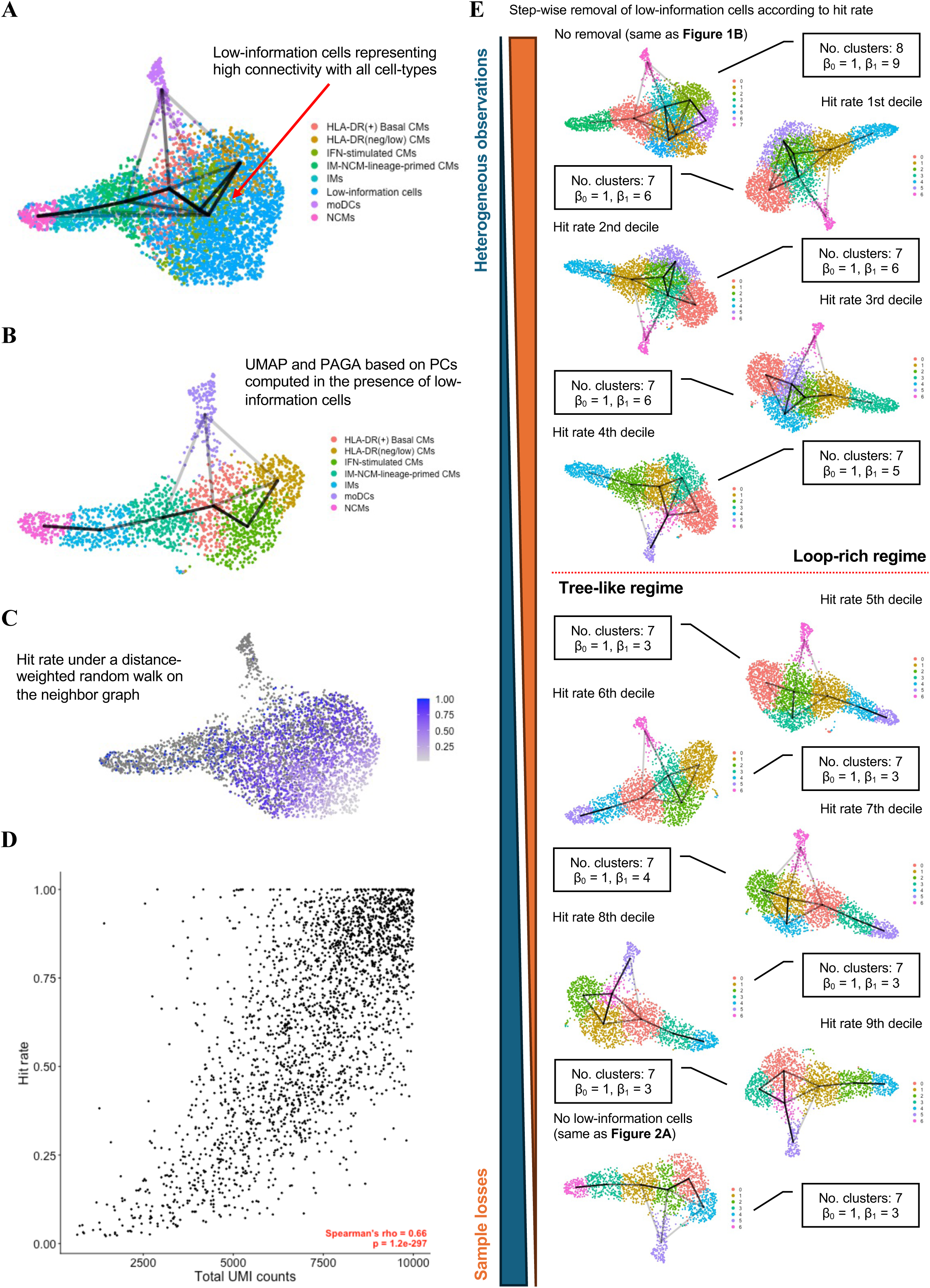
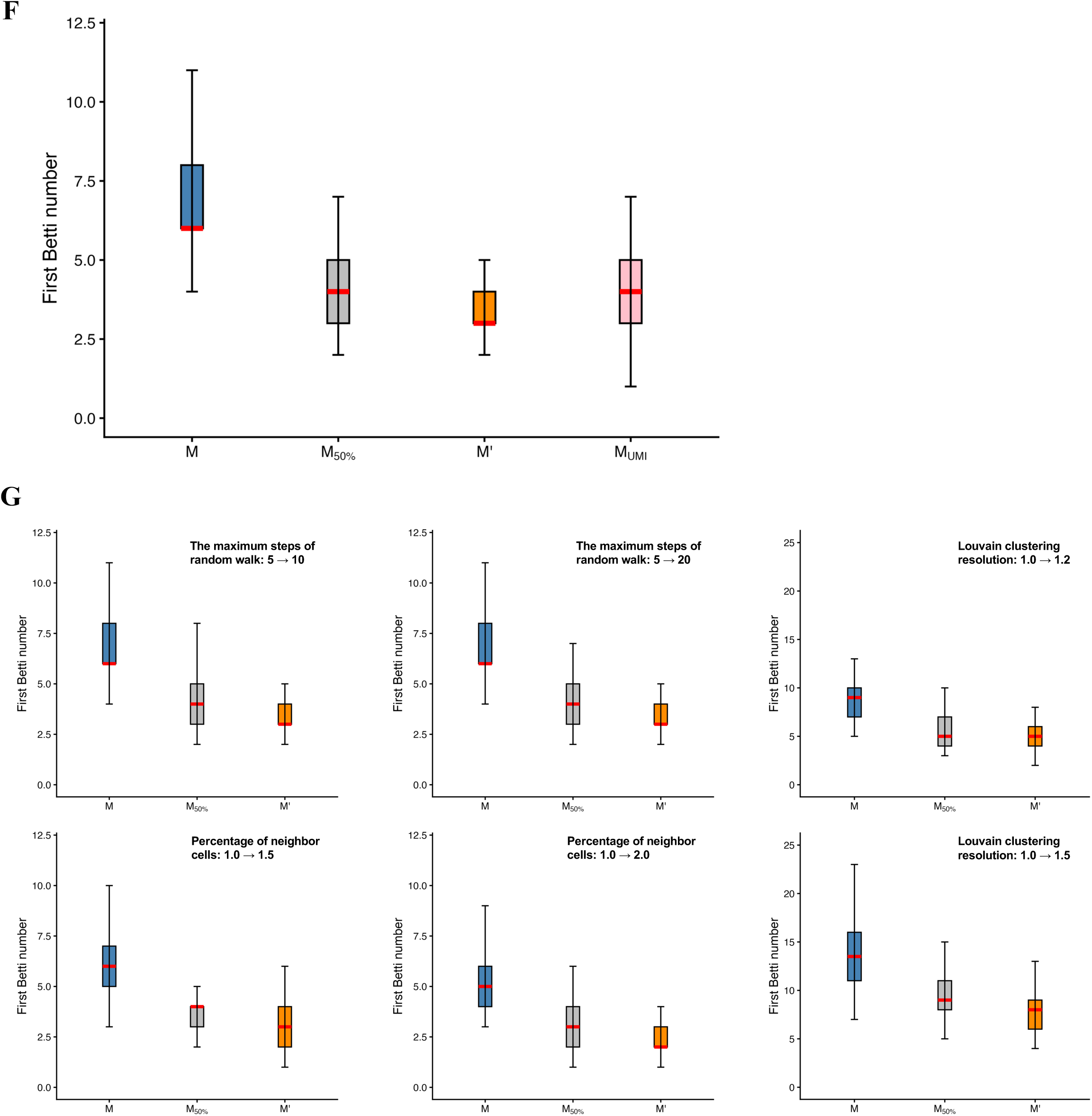
Topological descriptors of the data-manifold skeleton resolve the trade-off between preserving low-dimensional structure and minimizing sample loss. **A** UMAP embedding of *M* overlaid with PAGA edges representing connectivity between annotated clusters inferred from the analysis of high-information cell population *M*′ alone (HLA^neg/low^ CMs potentially immunosuppressive or exhausted, IFN-stimulated CMs, HLA^+^ basal CMs, IM-NCM-lineage-primed CMs, IMs, NCMs, moDCs) and low-information cells defined by total UMI counts < 10,000. Only edges with connectivity ≥ 0.05 are shown, and the intensity of each line segment reflects the strength of connectivity. Cluster 1 (red arrow), which is predominantly composed of shallowly-observed cells, shows strong connectivity with many other clusters; **B** UMAP embedding of *M*′ which is computed by PCs calculated in the presence of low-information cells overlaid with PAGA edges representing connectivity between the annotated clusters. Only edges with connectivity ≥ 0.05 are shown, and the intensity of each line segment reflects the strength of connectivity; **C** To quantitatively assess, among low-information cells, those located near the data manifold spanned by high-information cells, we computed a hit rate to high-information cells on the neighbor graph of *M* constructed by the workflow (#). For each low-information cell, we performed 1,000 *k*-step weighted random walks following the graph and calculated the proportion of walks that reached any high-information cell (hit rate). The figure shows a feature plot of the hit rate for each low-information cell at *k* = 5, with the color bar indicating the hit rate. High-information cells are shown as gray dots; **D** Scatter plot of hit rate versus total UMI counts for low-information cells. The x-axis shows total UMI counts and the y-axis shows the hit rate at *k* = 5; the monotonic association between the two variables was assessed using Spearman’s rank correlation (𝜌 and P value); **E** Changes in the topological properties of the low-dimensional skeleton as cells with decreasing reliability are progressively removed according to their hit rate. From left to right, UMAP embeddings and PAGA graphs (connectivity threshold = 0.05) are shown for populations with no removal (same condition as Figure 1E) and after removing the bottom 10%, 20%, 30%, …, and 100% of cells ranked by the hit rate (100% corresponding to the condition in Figure 2B). For each step, we report the number of clusters in the PAGA graph and the number of independent loops (first Betti number, 𝛽₁). The resolution parameter for Louvain clustering on the neighbor graph was fixed at 1, and the number of neighbors was set to 1% of all cells; all topological properties were compared under these conditions; **F** Boxplot of the 𝛽₁ distributions from 100 repeated PAGA-based manifold inferences, each performed on random subsamples of 1,500 cells drawn from *M*, *M*_50%_denoting the cell population obtained after filtering out 50% of the low-information cells from *M* by the hit rate, and *M*’. *M*_𝑢𝑚𝑖_ contains the same number of cells as in *M*_50%_ extracted from *M* in descending order of total UMI counts, and the results of repetitive 𝛽₁ evaluations performed on the subsamples from *M*_𝑢𝑚𝑖_ are also shown. Red bars indicate the median; **G** Boxplot of the 𝛽₁ distributions from the repetitive manifold inferences, each performed on random subsamples of 1,500 cells drawn from *M*, *M*_50%_ , and *M*’ with several hyperparameter settings (e.g., the maximum steps of random walk, the percentage of neighbor cells, Louvain clustering resolution). *M*_50%_ was calculated by hit rate under each hyperparameter setting. Red bars indicate the median.

Intuitively, if a low-information cell lies close, in terms of geodesic distance, to high-information cells, then a random walk starting from this cell should reach the high-information cell subset within a small number of steps. Conversely, cells that remain among low-information cell population for many steps are expected to reside in regions where low-information cells are spuriously connected to one another. To formalize this, we performed *k*-step random walks (*k* = 5) from each low-information cell along the weighted neighbor graph learned on *M* and, over many trials (1,000 per cell), computed the probability of hitting any high-information cell within *k* steps, which we termed the hit rate. Consequently, cells with low hit rates were confined to specific regions in UMAP, and though the hit rate was positively correlated with total UMI counts, at the same time, cells with comparable total UMI counts exhibited a wide range of hit rates (Figure 4C, D). These observations suggest that, while the hit rate partially depends on observation depth, it also reflects geodesic proximity to high-information cell population on the data manifold and can serve as a quantitative indicator of the reliability of each low-information cell.

Furthermore, we used the hit rate to order low-information cells to remove them progressively from *M*, and applied the workflow (#) and PAGA under the same rules governing certain percentage (e.g. 1%) of neighborhoods in whole population and certain resolution at Louvain clustering. As low-information cells were removed according to the hit rate, the first Betti number (𝛽₁), which counts independent loops in a low-dimensional manifold skeleton, decreased and then plateaued when approximately half of the low-information cells had been discarded, after which further removal no longer reduced the number of loops (Figure 4E).

Additionally, to compare manifold structures generated by different cell populations under identical conditions, we randomly subsampled the same number of cells (e.g., 1,500 cells) from *M* − *M*_50%_denoting the cell population obtained after filtering out 50% of the low-information cells from *M* by the hit rate − and *M*′, and quantified their manifold structures using PAGA; this procedure was repeated 100 times. Comparison of the 𝛽₁ distribution across the three cell populations showed that, relative to *M*, both *M*_50%_ and *M*′ exhibited distributions shifted toward lower 𝛽₁ (Figure 4F). Further, when the same number of cells (*M*_UmI_) as in *M*_50%_ were extracted from *M* in descending order of total UMI counts, followed by the same subsampling procedure and evaluation of the manifold skeleton using PAGA, the resulting distribution of 𝛽₁ was similar to that of *M*_50%_ . This finding suggests that, in cell thresholding based on graph topology, total UMI count may also function as a metric comparable to the hit rate.

Sensitivity analysis showed that the changes in 𝛽₁ across *M*, *M*_50%_, and *M*′ were generally maintained under changes in hyperparameter settings, including the number of random-walk steps, the number of neighbors, and the clustering resolution (Figure 4G). Although it should be noted that the range of 𝛽₁ can be directly influenced by these hyperparameter settings and should therefore be interpreted in relation to the settings used for the biological annotation of high-information cells, the ranking among *M* , *M*_50%_ , and *M*′ was preserved irrespective of the hyperparameter choices. These results suggest that topological descriptor-based thresholding of *M* can stabilize manifold inference while minimizing sample loss.

## Discussion

Changes in cellular state arise from numerous intercellular interactions and intracellular signaling events, and single-cell omics invite us to view each snapshot of a tissue not as a static picture, but as the aggregate outcome of them unfolding over time. Within a given cell lineage, if the repertoire of attainable molecular states is sampled densely enough, the neighbor graph should, in principle, encode which states can be reached from which others under biological constraints. This motivates a manifold perspective in which cells with high dimensional feature vectors are regarded as samples from a low-dimensional cell-state manifold, and intrinsic cell–cell interactions, together with external perturbations such as drug exposure where present, manifest as constrained trajectories. In recent years, methods such as Perturb-seq and single-cell foundation models have emerged as approaches for quantifying and predicting changes in cell states induced by interventions^24,25^. A central problem is therefore to reconstruct, from finite samples, the geometry and coarse topology of this manifold.

Graph abstraction of single-cell data such as PAGA provides a practical approach to this problem. Constructing a coarse-grained neighborhood graph between clusters can be interpreted as a low-dimensional skeleton of data manifold; in particular, the topology of the PAGA graph—its connected components and loops—summarizes the degrees of freedom in state transitions inferred from the data. Our results support the view that such graph-based representations can, when built under appropriate conditions, capture aspects of the constrained degrees of freedom in biology that have classically been described as differentiation paths. This is especially valuable in medicine, where longitudinal sampling of the same tissue over time is often ethically or practically infeasible and we are effectively forced to interpret snapshot data about the underlying dynamical system.

However, in biomedical omics the act of observation itself introduces a characteristic “ambiguity”^26^. In droplet-based scRNA-seq, one of the most prominent sources of ambiguity is the strong heterogeneity in per-cell amounts of observed molecules. As our computational simulations demonstrate, shallowly-observed cells tend to appear ambiguous, in that they resemble multiple well-resolved states that are clearly separable among deeply-observed cells. Such ambiguous cells act as *intermediate states* that create illusory bridges between clusters in graph abstraction. In our monocyte example, when we restricted the analyses to the homogeneous observations from deeply-observed cells alone, the PAGA graph formed several local structures consistent with prior experimental and clinical work indicating HLA upregulation relevant to IFN stimulation^11,14,15^ and differentiation toward IM-NCM lineage^12,13^ or moDC^10^ from CMs, and also showed the sequence of these local structures. By contrast, under heterogeneous observations, illusory loops generated by the spurious clusters made a low-dimensional manifold skeleton no longer clearly identifiable.

This simple consequence of heterogeneous observations that shallowly-observed cells tend to connect to one another and to multiple clusters in the neighborhood graph is not merely noise. This destabilizes the very notion of “local neighborhood” in the underlying metric space on which graph construction is based. This issue is even less negligible and more complex in the integration of multi-layered modalities with differing underlying observation depths. In our analyses, imputation methods that rely on feature-space neighborhoods therefore show only limited benefit under realistic heterogeneous observations: when, at least locally, neighborhoods are already unreliable, propagating information along them can reinforce, rather than repair, the distortion of manifold inference. Restricting attention to sufficiently homogeneous observations can partially mitigate this issue. Yet, if a dataset already contains enough high-information cells to support a reliable neighborhood graph and a stable topology of graph abstraction, then using their observed expression values directly may be more consistent with the long-standing experimental ethos of biology that values raw measurements. In practice, a safe strategy would be to first perform manifold inference using the workflow (#) on the unfiltered cell population, examine whether shallowly observed cells aggregate, and then define high-information cells according to the average observation depth of other clusters.

At the same time, our results do not imply that all structure in the data is hopelessly corrupted by the ambiguity due to heterogeneous observations. Prior work has shown that technical noise and *dropout* can substantially affect linear and non-linear compressions for low-dimensional embeddings^6,23,27–29^. Our analysis extends this view by showing that, even under realistic levels of observational ambiguity, certain global and topological aspects of the data manifold can remain relatively stable within subsets of cells that carry sufficient information (Figure 4A, B). Moreover, total UMI-based thresholding is practically useful because it can be implemented with low computational cost, and topological descriptors may also serve as a useful guide for determining an appropriate boundary. Using the topological descriptors of graph abstraction, we can explicitly flag regions where the inferred structure is likely driven by observational ambiguity rather than true dynamics, and consequently, carve out a trustworthy proxy for the underlying biological constraints. Furthermore, improving the validity of computational inference is also important for the design of subsequent biological validation. For instance, in animal models, cell populations corresponding to particular states along an inferred trajectory could be isolated and subjected to tracing experiments using fluorescent dyes or similar labeling strategies, allowing lineage relationships and the directionality of state transitions to be examined experimentally. Because mathematical inference inevitably requires biological validation, this need itself cannot be avoided. However, the experimental strategies and even the underlying conceptual framework may differ substantially depending on whether validation is conducted under the assumption of a loop-rich manifold skeleton or under a manifold skeleton that more closely accords with conventional biological understanding of lineage relationships. Therefore, improving the validity of computational inference is important not only for enhancing interpretability at the computational level, but also for ensuring the appropriateness of downstream validation workflows, which we consider to be a central aspect of computational biology.

Finally, we emphasize that these problems are not solely the responsibility of end-users of single-cell analysis pipelines. In interdisciplinary fields such as computational biology, methodological innovations are often packaged and disseminated as broadly applicable solutions, while the underlying geometric and statistical assumptions—and the trade-offs they entail—are communicated only partially^30,31^. Our findings suggest that, for methods that operate on manifold approximations and neighborhood graphs, developers should explicitly articulate not only what additional “information” their models appear to recover, but also which assumptions about distances, neighborhoods and dynamics are being relaxed or violated in the process. The framework we introduce in this work is, of course, not exempt from these considerations. It presupposes that, within a given dataset, there exists a sufficiently large subset of high-information cells to anchor a faithful reconstruction which can be proxies locally for at least part of the underlying biological constraints. Furthermore, our graph-topology-based sample thresholding should be applied after first classifying the overall cell population into highly distinct groups roughly, and then used when performing analyses of continuous lineages within each group. In this sense, it is less a recipe for extracting ever more information from single-cell data than a framework for deciding which biological narratives—namely, those concerning the degrees of freedom underlying differentiation and response to stimuli—we are prepared to trust.

## Methods

### Computational environment and package list

In computational simulations conducted in Python, core numerical and data-handling libraries included NumPy, pandas, and SciPy. Visualization was performed using Matplotlib. HDF5-based input/output was handled through h5py, and progress monitoring used tqdm.

All R-based analyses were performed in R 4.5.0. Core packages included Seurat and SeuratObject for single-cell analysis; tidyverse for data manipulation; plotly, scales, and htmlwidgets for visualization; hdf5r and SeuratDisk for HDF5-based I/O and object conversion; and SeuratWrappers for auxiliary single-cell workflows. zellkonverter was used for interoperability between Seurat objects and AnnData. Python-based steps were accessed from R via reticulate. Where applicable, imputation analyses used scImpute and SAVER. PAGA were executed from R using the reticulate interface. A dedicated conda environment named r-scanpy was created, and core dependencies were installed from conda-forge (including scanpy, anndata, NumPy, SciPy, pandas, Matplotlib, h5py, umap-learn, python-igraph, leidenalg, and pynndescent). All hyperparameters in the analysis were deposited on the corresponding author T.T. GitHub Page (https://github.com/Tomo-koko/Heterogeneous-single-cell-observation) and *Zenodo* with DOI: 10.5281/zenodo.17925918.

### Calculation of UMI distribution in empirical scRNA-seq datasets

Using the empirical UMI count matrix from real scRNA-seq datasets (in vivo, human: PBMC by 10x Genomics, 36,601 genes × 11,996 cells^7^; in vitro, human: lung progenitor organoids by Y. Miao *et al*, 36,601 genes × 14,100 cells^20^; mouse: complex sample containing cells from a tissue and cell lines by E. Mereu *et al*, 34,900 genes × 4,996 cells^8^), we calculated per-cell *N*(*k*), the number of genes with exactly *k* observed UMIs. We then stratified cells by their total UMI counts into bins of width 1,000 and computed *N̄*(*k*), the mean of *N*(*k*), for each bin. For each dataset, we used the publicly available filtered UMI count matrix. The source of each dataset is in the **Data Availability**.

### Manifold-based single cell analysis of empirical PBMC scRNA-seq data

In general, a workflow (#) has been used to analyze scRNA-seq data:

I. Log1p normalization of gene expression in each cell based on total UMI counts.
II. Identification of highly variable genes (HVGs).
III. Scaling of gene expression across cells (mean 0, variance 1).
IV. Principal component analysis (PCA).
V. Neighbor graph construction, clustering, and UMAP embedding in the PCA space.

We first extracted viable cells with %*M*𝑇 < 15, where denotes the percentage of UMIs mapping to mitochondrial genes, from the PBMC scRNA-seq UMI count matrix and applied the workflow (#) to obtain a coarse classification of cell types. Based on differential expression analysis between clusters, we identified clusters showing monocyte- like gene expression patterns, lymphocyte-like patterns, and other patterns (e.g., platelets).

We next re-applied the workflow (#) to the set of clusters representing monocyte characteristics to assess contamination by mRNA derived from other cell types, i.e., potential doublets. Differential expression analysis among the resulting clusters revealed that cell populations forming discrete “islands” on the UMAP showed evidence of lymphocyte-derived gene expression; these cells were therefore excluded as doublets. The remaining cells were defined as the putative monocyte population *M*. Up to this point, parameters such as the number of neighbors and the number of principal components in the standard analysis were chosen within commonly used ranges. For all subsequent analyses of the PBMC dataset, we basically adopted the following rule: among all observed cells, we constructed a weighted graph by connecting each cell to the top 1% of cells with the smallest Euclidean distance in a thirty-dimensional PCA space, and we interpreted the resulting Louvain modules using a fixed resolution parameter of 1. All the parameters used in each workflow (#) and slightly-modified pipelines for validations by the mouse scRNA-seq data were deposited in the corresponding author T.T.’s GitHub Page (https://github.com/Tomo-koko/Heterogeneous-single-cell-observation).

Next, we applied the workflow (#) to *M* to obtain a PCA embedding and a Euclidean distance–based neighborhood graph. We visualized the distribution of total UMI counts in each cluster using violin plots, and additionally generated three-dimensional dot plots in which the UMAP embedding provided the x–y coordinates and total UMI counts were mapped to the z-axis, in order to examine whether total UMI counts were unequally distributed within the UMAP- derived low-dimensional manifold. To evaluate inter-cluster connectivity when interpreting Louvain clusters as a coarse-grained partitioning of cell states on the manifold, we applied partition-based graph abstraction (PAGA)^9^ to *M*. The neighborhood graph used by PAGA was constructed with the same settings as in the workflow (#). Throughout this study, the PAGA connectivity threshold was fixed at 0.05; edges were drawn only between clusters whose connectivity exceeded this threshold, and the resulting graph was visualized as an overlay on the UMAP embedding and used for subsequent topological analyses.

### Reconstruction of data manifold using high-information cells in empirical PBMC scRNA-seq data

*M* is a cell population with heterogeneous observation depth, that is, a mixture of shallowly- and deeply-observed cells. Among these, we defined high-information cells as those observed at sufficiently high depth and analyzed the data manifold in *M*′, the subset consisting only of high-information cells. Note that the threshold defining *M*′ is determined relative to the UMI count distribution of the parent population *M* and is not guaranteed to serve as a generally applicable threshold across all datasets. In this study, we set a threshold of 10,000 total UMIs so as to include approximately the top 40% of cells, extracted high-information cells with total UMI counts > 10,000, and defined this subset as *M*′.

We then applied the workflow (#) to *M*′ and, as for *M*, constructed a neighborhood graph and performed clustering to obtain a coarse-grained partitioning of cell states on the data manifold. After confirming that the localized enrichment of shallowly-observed cells seen in the UMAP of *M* was no longer apparent, we assigned biological annotations to clusters based on differential expression analysis and the expression of representative marker gene sets (e.g., classical monocyte (CM) marker genes, non-classical monocyte (NCM) marker genes, HLA-related genes, interferon-stimulated genes (ISGs), and dendritic cell (DC) marker genes). Finally, using PAGA, we evaluated connectivity among the cell-state partitions obtained in *M*′ and examined whether the local structure of the low-dimensional skeleton of the data manifold reflects phenomena that are supported by known biology.

### Imputation and normalization in PBMC scRNA-seq analysis

In scRNA-seq analysis, observed zeros are often regarded as a mixture of technical zeros (e.g., failures of cDNA reverse transcription/amplification or shallow sequencing) and biological zeros (e.g., true absence of expression), collectively referred to as *dropout*. Numerous normalization and imputation methods have been developed to address such technical zeros using neighborhood graphs, and we employed representative methods—SCTransform^16^, ALRA^17^, SAVER^18^, and scImpute^19^—to test whether compensating for dropout can correct distortions of the data manifold caused by heterogeneous observation depth in scRNA-seq. UMI count data were used as input, and analyses were performed on the 4,410 cells contained in M. Parameters for each imputation method were set to their default values unless otherwise specified. Downstream analyses using the workflow (#) were then conducted on the normalized/imputed data generated by each method. For dot plots visualizing the three-dimensional distribution with UMAP coordinates on the x–y plane and total UMI counts on the z-axis, the z-axis corresponded to total UMI counts from the raw (pre-normalization/imputation) count data. For scImpute, we specified seven clusters, corresponding to the number of monocyte cell states that appeared appropriate based on analyses restricted to *M*’.

### Simulation of expression distribution models

We considered a typical human gene panel comprising 36,601 genes. For each simulated cell, we defined a cDNA pool derived from its mRNA molecules and assigned gene-specific cDNA fractions as follows. Let *p* denote the fraction of gene-derived cDNA in the pool, and order genes in ascending order of *p*, assigning gene IDs 𝑖 accordingly. The fraction for gene 𝑖 is denoted *p*_*i*_. It should be noted that the distribution of *p*_*i*_ corresponds to the intracellular gene regulatory network, and that gene IDs 𝑖 do not denote an existing gene identifier (e.g., a gene symbol), but simply specifies the rank order obtained by sorting *p* according to its magnitude.

We next considered how *p*_*i*_ increase as gene IDs increase; a linear increase model, an exponential increase model, and a combinational increase model of *p*_*i*_. Given that mRNA molecules, as the source of cDNA, are discrete physical entities with non-negative integer counts per gene and a total per cell on the order of 10^5^ molecules, we computed *p*_*i*_ under this constraint. The simulation was performed under the assumption that all UMIs are equally probable and that each captured molecule is labeled with a distinct UMI. Based on previous estimates that the total number of genes expressed in a single cell is ∼ 10,000, we further assumed that only genes with indices 𝑖 ≥ 26,602 can be expressed (i.e. *p*_*i*_ ≥ 0 for 𝑖 ≥ 26,602 and *p*_*i*_ = 0 otherwise)^21,22^. *p*_*i*_ were calculated as below under each model assumption:

- Linear increase model:

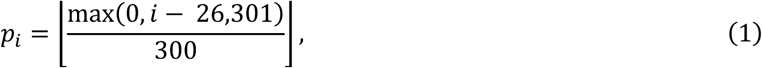

where ⌊𝑋⌋ denotes a maximum integer which does not exceed 𝑋.

- Exponential increase model:

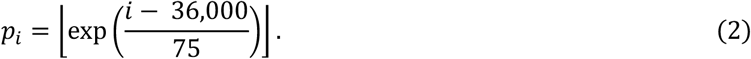

- Combinational increase model

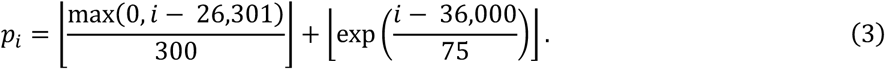

Under a Poisson sampling model for UMIs drawn from the cDNA pool, 𝑋_*i*_, the observed UMI count for gene 𝑖 and 𝔼[*N*(*k*)], the expected *N*(*k*) are given by these equations, respectively:

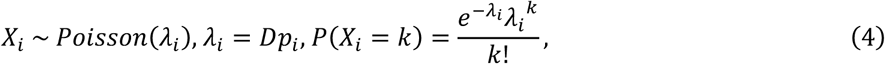

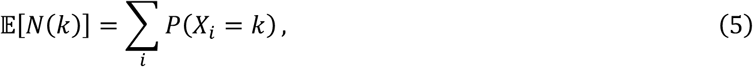

where 𝐷 denotes the total number of sampled UMIs.

To incorporate overdispersion assumed in scRNA-seq data, we also simulated 𝔼[*N*(*k*)] under a negative binomial (NB) sampling model:

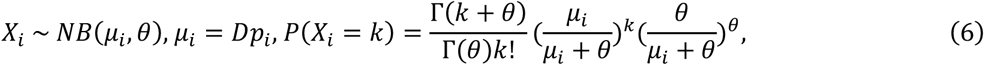

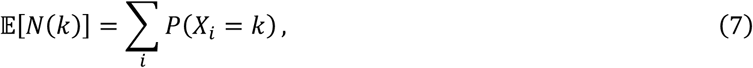

where *θ* is a dispersion parameter, and computed 𝔼[*N*(*k*)] for *θ* = 0.1, 0.3, 0.5, 1, 3, 10.

### Simulation modeling cell-type discrimination, cell-state discrimination, and step-wise transition

Building on the model (*), we generated a simulated cell population 𝑋 modeling cell-type discrimination as follows:

- Cell 1: 2,000 cells generated according to the model (*).
- Cell 2 and 3: 1,000 cells generated from a distribution obtained by randomly permuting the gene IDs of model (*) with different random seeds.

Building on the model (*), we next generated a simulated cell population 𝑌 modeling transcriptomic changes such as pathway activation as follows:

- Cell 1: 2,000 cells generated according to the model (*).
- Cell 4: 2,000 cells generated from Cell 1 by scaling the cDNA counts of 300 genes randomly selected from gene IDs 26,602 − 36,601 by either 2 or 0.5.

Building on the model (*), we further generated a simulated cell population *Z* modeling step-wise cell state transition such as pathway activation and epigenetic de-repression with an assumed lineage as follows:

Assumed lineage: A → B (pathway activation) → C (further pathway activation)→ D or E (epigenetic de-repression)

- Cell A: 1,000 cells generated according to the model (*).
- Cell B: 1,000 cells generated from Cell A by scaling the cDNA counts of 300 genes randomly selected from gene IDs 26,602 − 36,601 by either 1.67 or 0.6, modeling signaling activation.
- Cell C: 1,000 cells generated from Cell B by further scaling the same set of 300 genes by either 1.67 or 0.6.
- Cell D and E: 1,000 cells generated from Cell C by selecting 10 genes randomly from gene IDs 1 − 26,601 and increasing their cDNA molecule counts in the pool from 0 to 50 with different random seeds, modeling epigenetic de-repression.

For each simulated cell, the total UMI counts were randomly drawn as an integer from a normal distribution with mean 5,000 and standard deviation 1,500 , and clamped at a minimum of 500 . We analyzed the simulated populations by the workflow (#), the three-dimensional visualization, and PAGA.

### Topology-guided thresholding of data manifolds based on random walk metrics

When attempting to recover a data manifold that reflects biological constraints under heterogeneous observation depth, one inevitably faces the question of how to remove low-information cells. Simple thresholding by depth ignores biologically meaningful differences in transcriptional activity across cell types, so a quantitative metric is needed to evaluate low-information cells using the data manifold reconstructed from high-information cells as an anchor.

First, we transferred the annotations defined on the data manifold of *M*′ (e.g., HLA^+^ basal CMs) back to *M*, and labeled all cells that had been excluded from *M*′ as “low-information cells.” We then used PAGA to evaluate connectivity between each annotation and the low-information cells in the PCA space of *M* generated by workflow (#). Next, while keeping this PCA embedding fixed, we restricted the graph to high-information cells and again assessed inter-annotation connectivity using PAGA. These analyses confirmed that, in *M*, the low-dimensional skeleton generated by high-information cells is preserved even in the presence of low-information cells.

Next, to quantify how close each low-information cell lies to the data manifold spanned by high-information cells—that is, how easily it can *escape* the low-information cell population by following the neighbor graph—we considered *k*-step random walks on the weighted neighbor graph in PCA space, initiated from each low-information cell. We defined the hit rate as the proportion of random-walk trials in which a *k*-step walk starting from a low-information cell reached any high-information cell at least once. In this study, we set *k* = 5 and performed 1,000 trials per low-information cell to compute the hit rate for all low-information cells.

Finally, we identified low-information cells falling into the bottom 10%, 20%, …, 90% of the hit rate distribution, progressively removed these cells, and examined how the topological structure of the data manifold produced by workflow (#) and PAGA changed in terms of the number of clusters (nodes), the zeroth Betti number (𝛽_0_), and the first Betti number (𝛽_1_). By determining the hit rate threshold at which the behavior of 𝛽_1_shifts—separating a loop-dominated regime from one dominated by tree-like structures—we obtained a threshold for low-information cells that does not directly depend on depth but is instead grounded in the topological properties of the data manifold. The analysis was conducted using the following baseline hyperparameters: the number of neighbors was set to 1% of the total number of cells, the random walk length was set to 5 steps, and the Louvain clustering resolution was set to 1.0. As a sensitivity analysis, we then examined the behavior of 𝛽_1_ when these parameters were varied to 1.5% and 2.0% for the number of neighbors, 10 and 20 steps for the random walk length, and 1.2 and 1.5 for the Louvain clustering resolution.

To compare *M*, *M*_50%_ denoting the cell population obtained after filtering out 50% of the low-information cells from *M* by the hit rate, and *M*′ under the same conditions, we subsampled 1,500 cells randomly from each cell population and evaluated the behavior of 𝛽_1_by PAGA; this procedure was repeated 100 times. Because these were repeated subsampling from the same populations, independence was not preserved, and comparisons across groups were therefore made by visualizing the distributions with box plots.

### Research integrity including usage of GPT models

In this study, we primarily used the generative model ChatGPT as an auxiliary tool for portions of the R and Python coding and for drafting the manuscript. All code and text generated by the model were reviewed and verified by the corresponding author, and, to further safeguard research integrity, we screened the manuscript for textual similarity against internet-available literature using Turnitin. All schematic representations in this manuscript were created using BioRender and are published under an academic publishing license; the authors did not use generative AI for the creation of any schematic figures.

## Acknowledgements

This work was supported by Keio University Grant-in-Aid for Encouragement of Young Medical Scientists (to T.T.), Japan Society for the Promotion of Science (JSPS) KAKENHI Grant Number JP25K21344 (to T.I.), the Ishii-Ishibashi Foundation, and Japan Science and Technology Agency (JST) Grant Number JPMJPF2101. T.T. was supported by Takeda Science Foundation Scholarship, and several scholarships in Keio University Graduate School of Medicine. The authors thank to Ms. M. Abe and the members of Department of Extended Intelligence for Medicine for dedicated administrative supports. The author T.T. also thanks to Dr. M. Uchihara in Japan Institute for Health Security and Dr. K. Sekine and Mr. K. Maekawa in National Cancer Center Research Institute for useful advices.

## Authors Contribution

T.T. conceived and supervised the project. T.T. and Y.Y. designed the computational framework, performed the simulations and statistical analyses, and interpreted the results. Y.O. and T.I. performed validations of the results. T.T. drafted the manuscript, and Y.Y., Y.O., T.I., and K.S. contributed to revision and refinement of the text. All authors approved the final version of the manuscript.

## Data Availability

This study is based on previously published datasets. The human PBMC scRNA-seq dataset generated by 10x Genomics is available from 10x Genomics Web Page by Sample ID “10k_PBMC_3p_nextgem_Chromium_X”. The human lung progenitor organoid scRNA-seq dataset by Miao *et al*. is available from GSE250399. The mouse scRNA-seq dataset by E. Mereu *et al*. is available from GSE133535. No new sequencing data were generated in this study.

## Code Availability (including supplementary information)

The source codes and supplementary files related to this research were deposited in the corresponding author T.T.’s GitHub Page (https://github.com/Tomo-koko/Heterogeneous-single-cell-observation) and *Zenodo* with DOI: 10.5281/zenodo.17925918.

